# Global gene expression profile during low temperature in the brain of grass carp (*Ctenopharyngodon idellus*)

**DOI:** 10.1101/862102

**Authors:** Mijuan Shi, Qiangxiang Zhang, Yongming Li, Wanting Zhang, Lanjie Liao, Yingyin Cheng, Yanxin Jiang, Xiaoli Huang, You Duan, Lei Xia, Weidong Ye, Yaping Wang, Xiao-Qin Xia

**Author notes:** Correspondence authors: Xiao-Qin Xia and Yaping Wang.

## Abstract

Grass carp is an important commercial fish widely cultured in China. Large range of temperature, in particular extremely low temperature, has dramatic effects on the aquaculture of this teleost. However, there is relatively little research on the molecular responses in the fish exposed to cold. Given the limited vision of approaches targeting individual genes, we investigated the transcriptome profiles of brain in response to cold in order to comprehensively characterize molecular mechanisms behind it. This study indicated that the estrogen signaling pathway was inhibited in brain when grass carp acclimated to low temperature, while terpenoid backbone biosynthesis pathway and steroid biosynthesis pathway were significantly activated. Such a result implied the crucial role of cholesterol in cold acclimation. Moreover, plenty of differentially expressed genes associated with spliceosomes were enriched during cooling process, which suggested alternative splicing may be involved in the regulation of biological process in acclimation to temperature changes. In researches on extremely low-temperature tolerance, we identified four genes (DUSP1, HSPA6, NR4A1 and GADD45B) associated with MAPK signaling pathway. The four genes, extensively up-regulated at 4°C and remained relatively low expression at moderate temperature, were closely related with extremely cold condition. Further examination of the candidate genes can provide insights into the mechanisms of grass carp to endure extremely low temperature in the winter.

## 1. Introduction

As the “abiotic master factor” [1], water temperature controls all the physiological and behavioural parameters of fish [2]. In the past few decades, the world has suffered climate caprice which seriously endangered aquatic poikilotherms’ survival and damaged aquaculture industry [3]. Given the ecological and economic importance of fish, the influence of the interaction between fish and water temperature has been comprehensively studied but far from concluded.

While the effect of high-temperature on fish has been concerned and studied extensively due to the acceleration of global warming [4], low temperature caused much more fish death and brought acute economic losses to aquaculture industry [5]. Compared with studies on mammals and birds, the research on cold adaptation of fish is relatively limited and the underlying mechanisms remains obscure. Nevertheless, some researches have thrown light on the response of fish to cold [2, 6]. As a way to respond ambient temperature change, migratory fishes simply swim to suitable areas. However, non-migratory fishes have to tolerate low-temperature condition with special mechanisms. Indeed, fishes living in cold water areas with low-temperature throughout the year have unique physiological mechanisms to adapt the adverse conditions [7, 8]. The most notable example was the antifreeze protein in Antarctic fishes [9]. In addition, some studies revealed that the lack of hemoglobin and the synthesis of tubulin could improve the adaptation to low-temperature environments [10-12].

Meanwhile, eurythermic fishes, which adapt to a wide range of water temperatures, are supposed to have more complicate mechanisms to maintain homeostasis in response to fluctuation of ambient water temperature. Although many studies have paid attention to the effects of cold shock on fish, little exploration has been conducted on low-temperature tolerance. Limited researches have focused on the physiological phenomena and molecular mechanisms of fish organs or tissues at low temperatures, such as anion transport by mitochondrion-rich chloride cells in fish gills (Fundulus heteroclitus) [13], proteomic analysis of heart (Gillichthys mirabili) [14], changes of free amino acids in blood (Solea senegalensis), and the secretion of hormones in the brain [15, 16].

Especially, as the most important organ of the central nervous system, the brain not only plays crucial function in processing external information (including temperature), but also regulates homeostasis by inducing the secretion of hormones, such as growth hormone (GH), thyroid-stimulating hormone (TSH), and adrenocorticotropic hormone (ACTH). Throid hormone (TH) manipulated by TSH has been shown to be a regulator of heart and muscle function during cold acclimation [17-19]. Although the relationship between low-temperature and hormone secretion directly or indirectly manipulated by the brain has been demonstrated in fish, the mechanisms and functions of the biological process in low-temperature tolerance have not been fully elucidated. Moreover, the brain must maintain its normal physiological activities as well as mediating other organs of the organism. Previous researches implied that some compensation mechanisms allow for higher rates of metabolism in brain and increase brain fluidity at low temperature to maintain their functional and structural integrity during cold adaptation [20-22].

With the development of technology, the methods for analyzing the organismal, cellular and molecular responses of fish to temperature changes have made great progress [23-26]. Particularly, with the tremendous advances in high throughput sequencing technology, RNAseq, an important omics technique, has been playing a vital role in studying the systematic processes by which fish respond to environmental changes [27, 28]. Given the enormous economic value of grass carp and their special winter dormancy, we profiled the transcriptome in the brain during cold acclimation. Even though the process of adjusting the cold environment was gradual and continuous, the grass carp exhibited significantly different characteristics with the water temperature above or below 10°C. Hence, we divided the physiological process into two stages. The first stage is at a temperature going down from 27°C to 10°C, and the second stage is a temperature from 10°C to 4°C. We supposed grass carp had different mechanisms in cold adaptation process of continuous cooling and the low-temperature tolerance process initiated by low temperatures below 10°C. In this study, we primarily aimed to identify genes induced by low temperature in the brain, and to investigate the mechanisms of cold adaptation and low temperature tolerance of this eurythermic fish.

## 2. Materials and Methods

### 2.1. Fish and cold acclimation

Fifty 14-month-old grass carp, each one weighted around 100g, were maintained at 27±1°C in a 300L tank for 2 weeks, fed 3 times a day. The water temperature was gradually reduced from 27°C to 12°C at a rate of 0.25°C/h and maintained for 24h. Then the temperature was increasingly lowered to 4°C as the same method. After the cooling process, the water was incrementally heated to 12°C and ultimately to 27°C at the same rate of 0.25°C/h. At each point, the brains of eight randomly selected grass carp were obtained and grinded in liquid nitrogen. Then, 100mg grinded tissue was added 1ml Trizol reagent and stored at -80°C. The experimental samples used for RT-qPCR were treated with the same project in the preparation of RNA-seq samples. At each temperature piont, 3 fish were randomly selected and the total brain RNA of each fish was separately obtained for RT-qPCR verification. The use of all experimental grass carp was approved by the Animal Research and Ethics Committees of the Institute of Hydrobiology, Chinese Academy of Sciences.

### 2.2. RNA extraction and sequencing

Total RNA was extracted using Trizol according to a standard protocol (Life tech, USA). After the quality inspection, mRNA was enriched using oligo(dT) magnetic beads and fragmented with ultrasound. The double strands cDNA was synthesize using the mRNA fragments as template and purified. Finally, sequencing adaptors were ligated to the cDNA ends. The desired fragments were purified and enriched by 10-cycle PCR amplification. The library quality was checked by Bioanalyzer 2100 (Agilent). The qualified library products were used for single-ended sequencing via Illumina HiseqTM2000. The accession number of the raw data is CRA001859 (BIG: http://bigd.big.ac.cn/gsa).

### 2.3. Data processing and basic statistical analysis

NGSQCToolkit (version 2.3.3) [56] was used to eliminate non-useful data with default parameters. The remaining clean reads were mapped to the reference genome [29] by Tophat2 (version 2.0.7) [57] and the transcripts were assembled with StringTie (version v1.3.1c) [58]. The sequences were Blast to nr database and the ones with an e-value ≤1e-5 were annotated. The prediction of non coding RNA combined the usage of CPC2 (version 1.2.2) [59] and CPAT (version 0.1) [60].The reads counts and TPM of transcripts were calculated by the salmon (version 0.12.0) [61] with non-alignment algorithm. The default parameters were used in all softwares above. The results above were all recoded in the supplement 2.

### 2.4. Identification of differentially expressed genes and transcripts (DEGs / DETs) and enrichment analysis of pathways

The edgeR [62] was used for differential expression analysis of genes and transcripts through glm approach with the likelihood ratio tests [63]. The genes, which stably expressed with less than 0.2 coefficient of variation during the whole process, were used to estimate variance of samples and to identify DEGs. The criteria to define differential expression was false discovery rate(FDR) ≤0.01 and absolute log2 of TPM ratio ≥ 1 (ref). The pathways database was downloaded with the API interface provided by KEGG. Total 318 pathway maps, of which the ratio of grass carp genes annotated to all genes of the map was more than 20%, were selected for the enrichment analysis. The statistical test for enrichment analysis was performed by R scripts executing Fisher’s exact test with the p-value ≤0.05.

### 2.5. Validation of expression profiles using RT-qPCR

Given the instability of beta-actin expression in cold condition, the gene RPL13A was took as the internal control which steadily expressed during the cold adaptation [28]. Then 4 genes (5 transcripts) were selected and the length of PCR products was about 120-250bp (Primer list in Table S4) and Tm ∼55°C.

## 3. Results

### 3.1. The transcriptome assembly and statistic

We conducted the project that decreased the water temperature from 27°C to 4°C and increased it back to 27°C to simulate the temperature changing in nature. The RNA-seq data sets at five temperature points (27°C - 12°C - 4°C - 12°C - 27°C) which sequentially reached during process were obtained and named as A, B, C, D, E. Each data set has 8.45±0.26M clean reads with 96.03±0.76% mapping rate (Table S1). Subsequently, we implemented the transcriptome assembly using published gene structure annotation information of grass carp as a reference [29] and generated a new annotation file containing 52,580 transcripts of 37,531 genes.

With the reference genome information, all assembled transcripts were classified into twelve categories (Table 1). Apart from 32,811 reference transcripts, we found another 19,769 new transcripts which belong to eleven classes according to the structural differences with reference genes. Afterwards we predicted 9,747 non-coding RNAs (ncRNAs) containing 5,745 single exon transcripts (SETs), which accounted for 2/3 of the total SETs. After removal of predicted ncRNAs and unexpressed transcripts, the remaining 38,942 transcripts were used for subsequent analysis.

**Table 1.** The Statistics of the Transcript Classification.

| Class code |  | NO. | SET* | ncRNA* |
| --- | --- | --- | --- | --- |
| = |  | 32811 | 3415 | 2903 |
| new<br>transcripts | j | 10088 | 0 | 1125 |
|  | u | 3605 | 2488 | 2869 |
|  | k | 2036 | 0 | 41 |
|  | p | 2035 | 2035 | 2005 |
|  | n | 842 | 171 | 156 |
|  | m | 828 | 106 | 81 |
|  | o | 597 | 0 | 174 |
|  | i | 329 | 286 | 306 |
|  | x | 94 | 2 | 69 |
|  | y | 22 | 0 | 17 |
|  | e | 1 | 1 | 1 |

SET: single exon transcript
ncRNA: non coding RNA

### 3.2. Differential expression of genes and transcripts

In order to investigate the consecutive and phased changes of the transcriptome profiles with the temperature fluctuating, we identified cumulative differentially expressed genes (cDEGs) / transcripts (cDETs) and phased differentially expressed genes (pDEGs) / transcripts (pDETs).

On one hand, we sequentially compared test groups B, C, D, E with control A to identify cDEGs and cDETs. As the temperature dropped from 27°C to 12°C, a total of 1,863 genes were differentially expressed with 4,875 cDETs. When temperature was continuously dropped to 4°C, the number of cDEGs and cDETs increased to 2,951 and 6,244. Conversely, when the water temperature was raised from 4°C to 12°C, the number of cDEGs and cDETs was slightly reduced to 2,782 and 6,134. In addition, as the water temperature rose to 27°C, we observed a sharp drop in the number of cDEGs and cDETs to 183 and 1,965 (Figure 1A).

**Fig 1.**
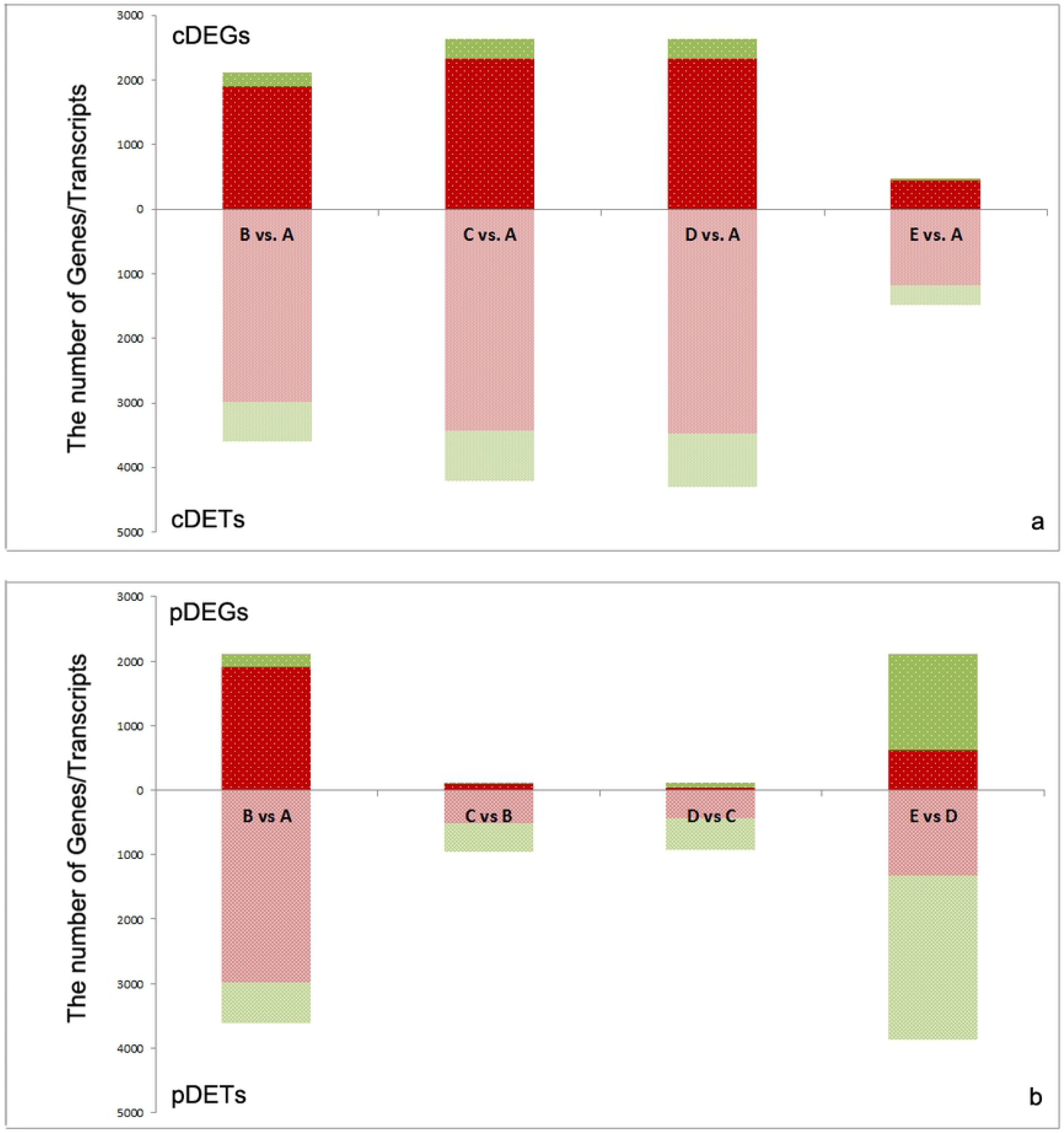
The stacked bar of DEGs and DETs. a. The cumulative differential expression genes (DEGs) and transcripts (DETs) comparing test group B (12°C), group C (4°C), group D (12°C) and gruop E (27°C) with control group A (27°C). b. The phased DEGs and DETs from comparisons of group B (12°C) vs. group A (27°C), group C (4°C) vs. group B (12°C), group D (12°C) vs. group C (4°C) and gruop E (27°C) vs. group D (12°C).In both two plots, red represents upregulated genes/transcripts and green represents downregualtion.

On the other hand, we separately investigated the difference of transcriptome profiles between every two adjacent temperature points (B vs. A, C vs. B, D vs. C, E vs. D) to find pDEGs and pDETs. Interestingly, as shown in Figure 1B, pDEGs and pDETs in C vs. B and D vs. C were considerable less than which of B vs. A and E vs. D. The number of pDEGs and pDETs were 84 and 1,716 in C vs. B, 114 and 1,770 in D vs. C, 1,863 and 4,875 in B vs. A, 2,551 and 5,888 in E vs. D.

### 3.3. The pathway analysis of cDEGs

KEGG analyses were performed to determine the pathways involved in responding to cold adaptation. Our aim was to ascertain the continuous changes in pathways in brain during cooling procedure. We separately enriched pathways with cDEGs from comparative analysis of four test groups and the control group (B vs. A, C vs. A, D vs. A and E vs. A). A total of 21 pathways were significantly enriched in metabolism, transcription, translation, signal transduction and cellular processes. In addition, we performed cluster analyses on the pathways enriched in the four sets of analyses in order to illuminate the dynamics of pathway combinations throughout temperature variation (Figure 2 a1). In oder to analyze the potential cross talk between different pathways, we constructed a network of enriched pathways using the common cDEGs as linkages (Figure 2 b1).

**Fig 2.**
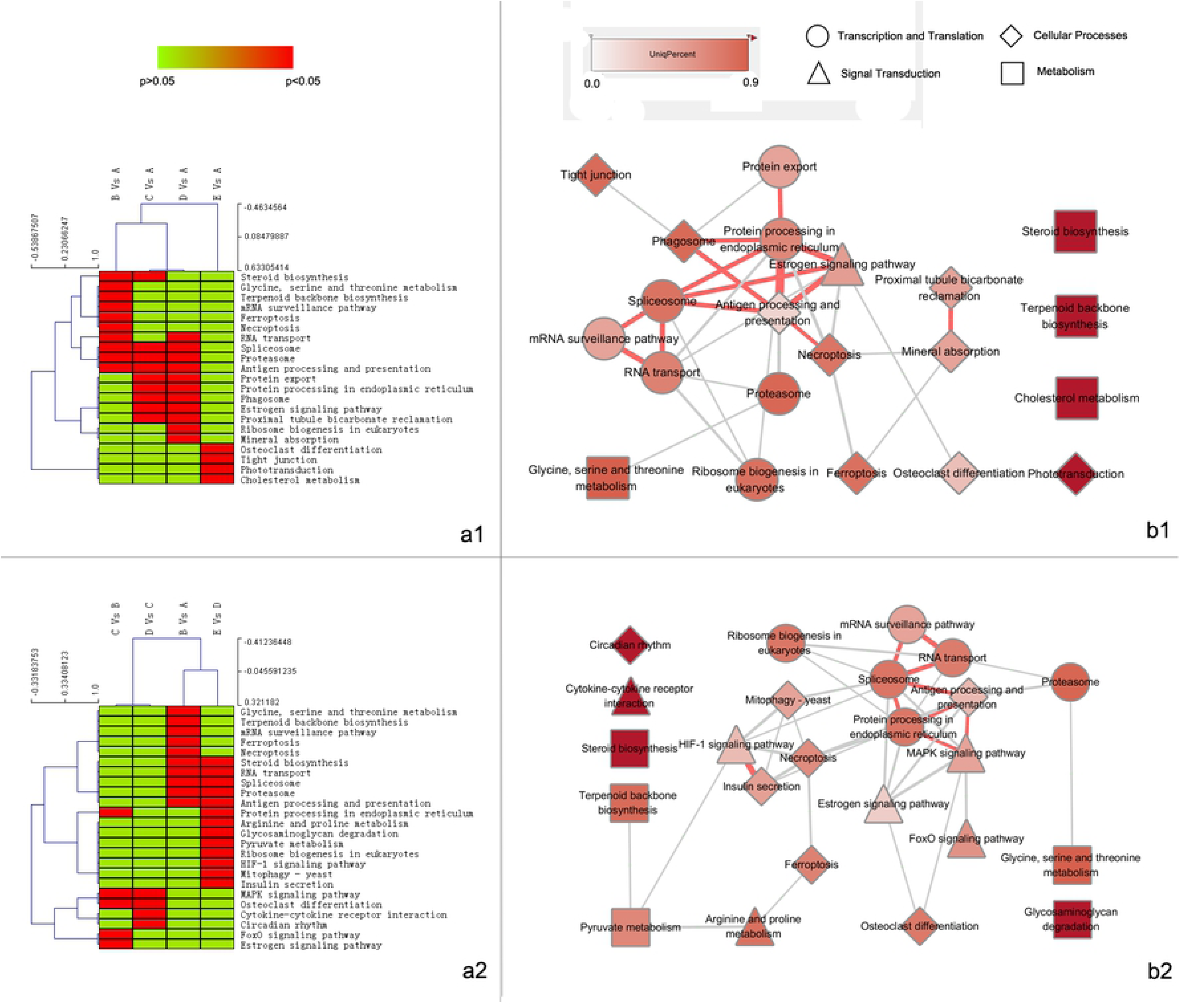
The heatmap and network diagram of enriched pathways. a1/a2. The heatmap of enriched pathways from cDEGs/pDEGs analysis. In each group, red represents the pathway is significantly enriched, while green is not. Clustering is performed based on the patterns of pathways profiling. b1/b2. The network of eriched pathways from cDEGs/pDEGs analysis. The pathways are classified into 4 categories: Transcription and Translation (Circle),Signal Transduction (Triangle),Cellular Processes (Diamond) and Metabolism (Square).The color depth of nodes represents the ratio of unique DEGs in the pathway. The width of edges represents the number of common DEGs shared by two adjacent pathways and the edges are marked with pink with more than 4 shared DEGs.

In the first stage with temperature decreasing from 27°C to 12°C, overall 10 pathways were significantly enriched including three for metabolism (glycine, serine and threonine metabolism, steroid biosynthesis, terpenoid backbone biosynthesis), two for apoptosis (necroptosis, ferroptosis), two for translation (RNA transport, mRNA surveillance pathway), one for transcription (Spliceosome), one for protein metabolism (proteasome), and one for immune system (antigen processing and presentation).

In the second stage of cooling with temperature down to 4°C, we found 9 conspicuous pathways: estrogen signaling pathway, steroid biosynthesis, spliceosome, phagosome, antigen processing and presentation, proximal tubule bicarbonate reclamation, and three pathways for protein metabolism (proteasome, protein export, protein processing in endoplasmic reticulum).

After stayed one day in 4°C, the water was slowly heated to 12°C. Although the pathway of steroid biosynthesis disappeared as well as RNA transport, two new pathways, mineral absorption and ribosome biogenesis, emerged in this phase, while the other pathways enriched in the second stage still remained significant in the third stage. As expected, all pathways identified in the low temperature stages were not significantly enriched when temperature went back to 27°C.

### 3.4. The pathway analysis of pDEGs

To eliminate the impacts of gene cumulative expression on pathway enrichment analysis, we investigated the phased changes of pathways using pDEGs between every two adjacent temperature points. The cluster analysis was subsequently implemented with the data from pathway analyses and the network of pathways was constructed using the same method as pathway analysis of cDEGs (Figure 2 b2). Given the similarity of associated pathways, comparisons between higher temperatures (B vs. A and E vs. D) were classed into one branch while comparisons between lower temperatures (C vs. B and D vs. C) belonged to the other one (Figure 2 a2). Intriguingly, pathways activated in moderate cold condition (12°C) were distinct from which in radical low temperature (4°C). We further paid attention to the MAPK signaling pathway, in which the associated pDEGs were highly sensitive to very low temperature, as shown by the gene expression heat map (Figure 3).

**Fig 3.**
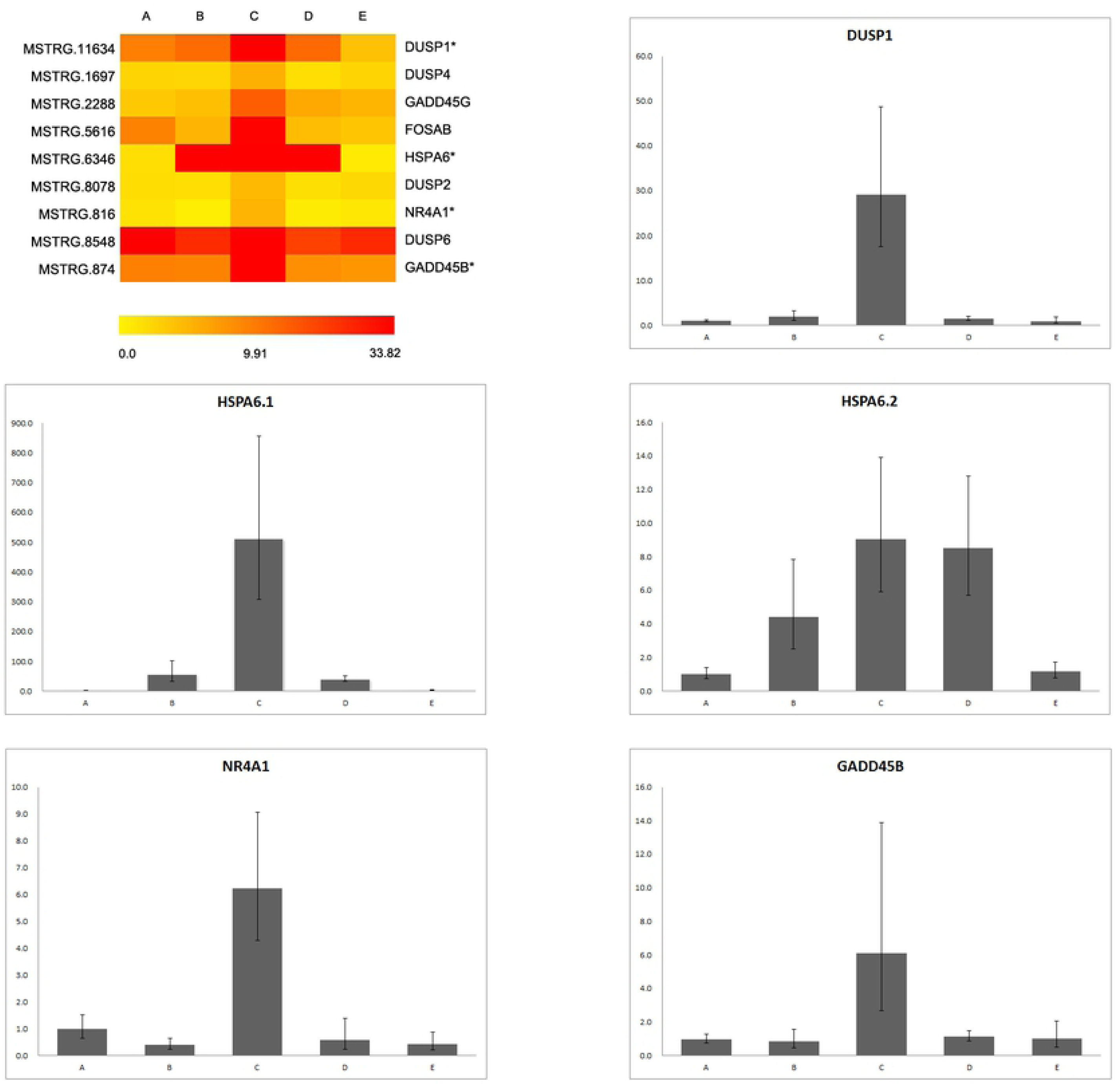
The expression profiles of pDEGs associated with MAKP signaling pathway. a. The heatmap of differential expression genes with the TPM in RNA-seq data. b1-b5. The expression profiles of DUSP1, HSPA6 (HSPA6.1, HSPA6.2), NR4A1 and GADD45B with the RT-qPCR results.

### 3.5. Validation of RNA-seq data with RT-qPCR

Considering the significance of MAPK signaling pathway in the cold response, we constructed a heat map of pDEGs associated with MAPK signaling using the RNA-seq data at five temperature points. Then four extensively highly expressed genes (DUSP1, HSPA6, NR4A1 and GADD45B) at 4°C to be verified by real-time quantitative PCR (RT-qPCR). Among these genes, two isoforms of HSPA6 (HSPA6.1 and HSPA6.2) were validated separately. The results of RT-qPCR were almost identical to those of RNAseq (Figure 3).

## 4. Discussion

The cold adaptation of fish was known as a continuous process which may last from days to months with the transcriptome consecutively changing [30]. However, at very low temperature in winter, many eurythermic fishes are able to enter dormancy, a dramatically distinct physiological status[31]. The obviously different response to moderate cold condition and extremely low temperature implies dissimilar mechanisms for cold adaptation. Grass carp enters dormancy when the ambient temperature is below 10°C. Therefore, we assume that grass carp have different mechanisms for cold acclimation during moderate cold and extremely cold periods. In our research, we attempted to illuminate the cold adaptation mechanism which functions throughout low temperature period and the low-temperature tolerance mechanism which activates during winter dormancy.

### 4.1 Mechanisms of cold adaption

Given the cascade amplification of signal transduction and hysteresis of gene expression, the changes of cumulative expression should be more representative of genes’ influence on biological processes. Hence, the pathway analyses of cDEGs could provide some clues for us to understand the underlying mechanisms of fish to adapt cold environment. As the clustering results revealed, we surveyed the emergence and persistence of pathways in different temperature stages during the cooling procedure.

#### 4.1.1 Hormones regulation

As many researches have confirmed that hormonal regulation played an important role in response to low temperature, the estrogen signaling pathway was significantly enriched in the second and third stages, and relevant gene expressions were down-regulated. This fact implies that the inhibition of hormones might be a crucial step in cold adaptation at very low temperature.

As an important class of hormones, estrogens participate in many physiological processes by binding to specific receptors [32]. Although there is relatively little evidence to unveil the relationship between estrogen and cold acclimation of fish, the correlation between inhibition of estrogen signaling pathway and the low levels of circulating vitellogenin in the plasma of male carp in winter was unfolded [33]. Additionally, many studies demonstrated the significance of estrogens in reproductive behaviors of fish [34, 35]. In the present research, we found the estrogen signaling pathway was considerably suppressed with a series of vital genes down regulated in 4°C and the inhibition still sustained with temperature rising to 12°C. These evidences may imply that suspension of reproductive behaviors of grass carp in low temperature condition was associated with the repression of estrogen signaling pathway in favor of individual survival.

#### 4.1.2 Lipid metabolism

Homeoviscous adaptation, an adaptive response of poikilothermic organisms in the cold, has been extensively studied with particular attention to the composition of membrane lipids [36]. Although the most observed alterations of membrane lipids involve a change of unsaturation of the fatty acids bound to phospholipids, the change of phospholipid to cholesterol ratio was confirmed to participate in the homeoviscous adaptation [37]. Recent studies have shown cholesterol synthesis probably involved in the cold acclimation of common carp [38] and yellow drum [39].

When water temperature decreased to 12°C in our research, the terpenoid backbone biosynthesis pathway was significantly enriched with six up-regulated cDEGs (HMGCR, HMGCS1, ACAT2, MVD, IDI1 and MVK). Meanwhile, eleven cDEGs associated with the steroid biosynthesis pathway were remarkably up-regulated, genes (ARHGAP32, CYP51, DHCR7, EBP, FDFT1, LSS, MSMO1, NSDHL and SQLEA) up-regulated at 12°C and genes (ARHGAP32, CYP24, DHCR7, EBP, FDFT1, LIPA, MSMO1, NSDHL and SQLEA) up-regulated at 4°C. Interestingly, the terpenoid backbone biosynthesis pathway is located the upstream of Steroid biosynthesis pathway. Combining the two pathways, we clearly observed an unimpeded route transforming Acetyl-CoA to cholesterol (Figure 4). Given the effects of cholesterol on manipulating fluidity and flexibility of cell membranes in the low temperature, we inferred the terpenoid backbone biosynthesis and steroid biosynthesis pathway played an vital role in grass carp to endure low temperature.

**Fig 4.**
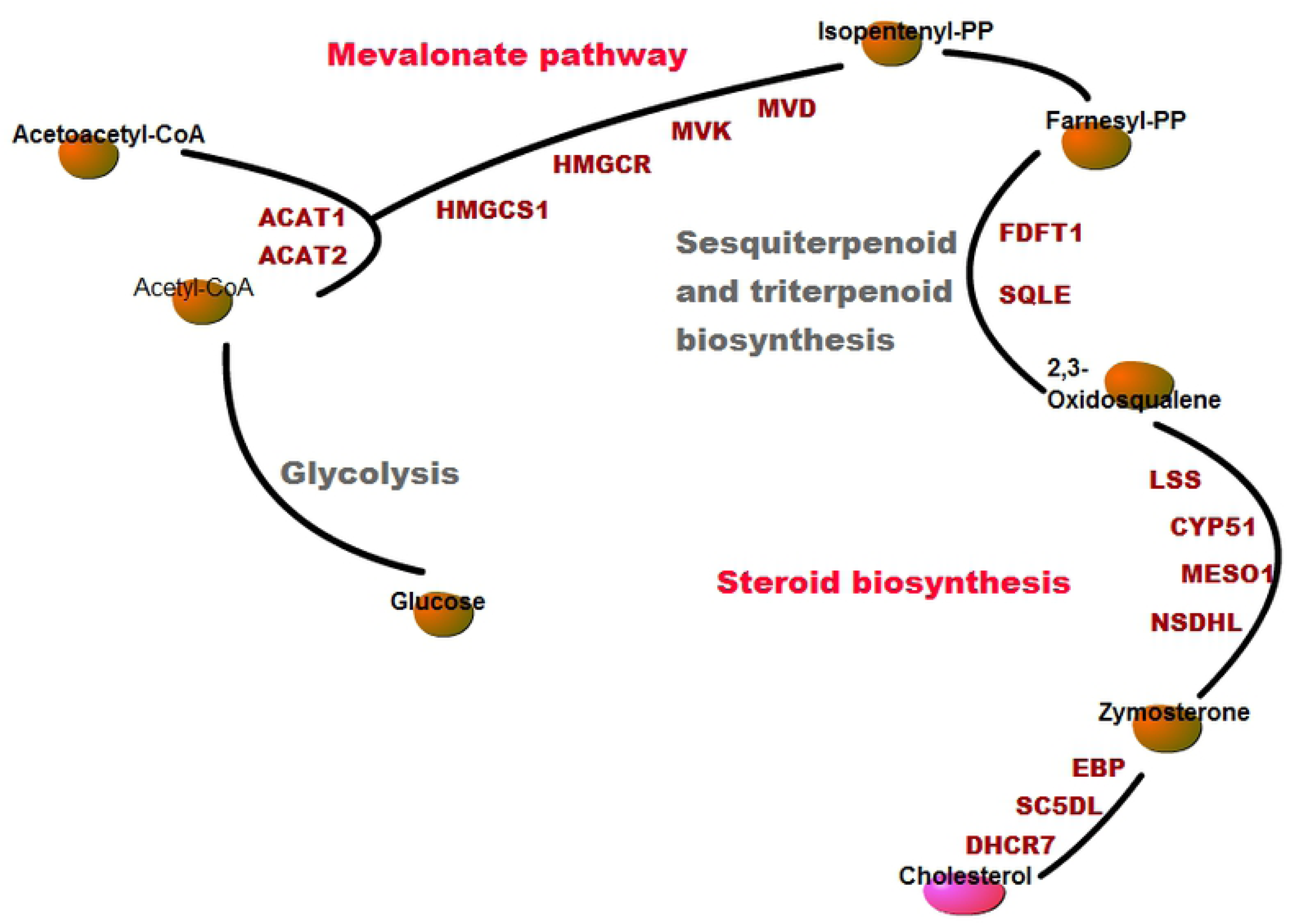
The schematic diagram of Cholesterol synthesis pathway.

#### 4.1.3 Alternative splice

Although the quantitative changes of gene expression was comprehensively and extensively explored on cold acclimation of organisms, relatively few studies have focused on the alternative splicing of genes. As an important post-transcriptional modification mechanism, alternative splicing can produce different transcripts from the same pre-mRNA. As the increasing studies on the role of alternative splicing in plant cold resistance are conducted [40, 41], some researchers have paid attentions on the effect of this mechanism on cold adaptation of fish. Alternatively spliced isoforms of delta 9-acyl-CoA desaturase distinctly respond to cold in common carp [42]. With RNA-seq analysis, alternative splicing of 197 genes were found to be modulated by cold stress in larval zebrafish [43]. Moreover, the comparative investigations on the extent of alternative splicing of genes in Atlantic killifish, threespine stickleback and zebrafish to respond cold indicated the universality and specificity of this mechanism in cold adaptation of fishes [44].

Similarly, our results revealed many transcript isoforms were sensitive to cold and have distinct expressional patterns from each other. Besides, we found the spliceosome significantly enriched throughout whole low-temperature phase. Therefore, we speculated that this post-transcriptional regulation was involved in the process of adaptation to cold for grass carp.

### 4.2 Mechanisms of low-temperature tolerance

While the cold adaptation of fish is a chronic and gradual process, extreme cold certainly bring more stress to organisms than moderate cold conditions. Clearly, the fact that the capability to survive from radically low temperature is crucial to fishes implies extremely low temperature possibly triggers unique mechanisms. In order to reveal the mechanisms of low-temperature tolerance in grass carp, we focused on the differentially expressed genes at 4°C compared to 12°C.

Surprisingly, the pDEGs (84) between 12°C and 4°C was dramatically less than those (1,863) between 27°C and 12°C. Of the 84 pDEGs, nine pDEGs (DUSP1, GADD45G, FOSAB, HSPA6, DUSP2, NR4A1, DUSP6, GADD45B, HSPA6.2) associated with MAPK signaling pathway were upregulated at 4°C.

#### 4.2.1 MAPK signaling pathway in response to extremely low-temperature

In general, MAPKs directly regulate the protein function through phosphorylation or indirectly influence the biological process by signal transduction. This pathway has been extensively concerned in freeze-resistance of plants [45, 46], and has recently been enriched in the liver proteomic analysis of *Takifugu fasciatus* during the cold acclimation [47]. Although the MAPK signaling pathway is involved in the adaptation to low temperature in plants and animals, the genes involved are different. Although the specific mechanism of MAPK signaling pathway in cold acclimation is unclear, the correlation between relevant genes and cold endurance of organism have been increasingly investigated. In present research, we focused on genes associated with MAPK signaling pathway and analyzed their expression patterns in response to temperature changes. Furthermore, we verified their expression profile by RT-qPCR.

Dual specificity phosphatase 1 (DUSP1) is a phosphatase which specifically phosphorylate tyrosine and threonine, and its expression increased in our RNA-seq data at 4°C, consistent with RT-qPCR reconfirmation. Knocking down DUSP1 can significantly increase the apoptotic rate of zebrafish ZF4 cells in low temperature [48]. This gene was also found highly related with low-temperature-induced embryonic diapause in Blue-breasted quail [49].

Nuclear receptor subfamily 4 group A member 1 (NR4A1), a member of the steroid-thyroid hormone-retinoid receptor superfamily, was confirmed to be upregulated in brown adipose tissue (BAT) of mice exposed to cold [50]. Relatively few studies reported the effect of NR4A1 in the social behavior in zebrafish [51, 52], and no report has been publicated about the relationship between NR4A1 and cold adaptation of fish. However, we found NR4A1 significantly up-regulated at 4°C in grass carp brain, while its expression remained in low-level at higher temperatures. The notable expression pattern of NR4A1 implied an important role in grass carp to tolerate the extremely low temperature.

Heat shock 70kDa protein 6 gene (HSPA6) is the 6th member of heat shock protein family A (Hsp70), a famous molecular chaperones family. The members of Hsp70 were often found active in the studies of cold acclimation or cold shock [53, 54]. Likewise, we found that HSPA6 expression increased significantly as the water temperature decreased, and the two isoforms of HSPA6 presented different expression patterns. One isoform (HSPA6.1) was highly up-regulated at 4°C, while the other isoform (HSPA6.2) incrementally expressed with temperature cooling from 27°C to 4°C. In consideration of the dissimilar sensitivity to temperature, we suppose that the two isoforms of HSPA6 have different impacts on the cold acclimation of grass carp.

Growth arrest and DNA-damage-inducible (GADD45) is a group of genes including three paralogs: GADD45A, GADD45B and GADD45G. Many studies have proven the involvement of GADD45 in the regulation of growth and apoptosis. Although a recent study revealed that the mice lacking GADD45G had defects in the thermogenic response to cold [55], we still know very little about the role of GADD45 in cold stress. According our results, both GADD45B and GADD45G seemed to be involved in this process, while the expression of GADD45A was not sensitive to temperature changes.

In the research, the response of grass carp to low temperature was divided into the cold adaption and the cold dormancy. The former was a systematic process, involving the hormone regulation, lipid metabolism, especially the cholesterol synthesis and the alternative splicing, as well as many other physiological changes and signal transduction. The genes involved in cold dormancy were enriched in the MAPK signaling pathway, which can directly regulate the protein function through phosphorylation.

## Author Contributions

Conceptualization, Y.W. and X.Q.X.; methodology, M.S.; software, W.Z. and M.S.; Validation, Q.Z., X.H., Y.J., Y.D., L.X. Y.C. and W.Y.; resources, Y.L. and L.J.; data curation and visualization, M.S.; writing – original draft preparation, M.S.; writing-review and editing, X.Q.X.; supervision, Y.C.; project administration, M.S.; funding acquisition, X.Q.X. and M.S.

## References

1. Brett, J.R., Energetic responses of salmon to temperature - study of some thermal relations in physiology and freshwater ecology of sockeye salmon (*oncorhynchus-nerka*). American Zoologist, 1971. 11(1): p. 99.

2. Donaldson, M.R., et al., Cold shock and fish. Journal of Fish Biology, 2008. 73(7): p. 1491–1530.

3. Brander, K.M., Global fish production and climate change. Proceedings of the National Academy of Sciences of the United States of America, 2007. 104(50): p. 19709–19714.

4. Logan, C.A. and B.A. Buckley, Transcriptomic responses to environmental temperature in eurythermal and stenothermal fishes. Journal of Experimental Biology, 2015. 218(12): p. 1915–1924.

5. Stauffer, B.A., et al., An oceanographic, meteorological, and biological ‘perfect storm’ yields a massive fish kill. Marine Ecology Progress Series, 2012. 468: p. 231–243.

6. Hu, P., et al., Global identification of the genetic networks and cis-regulatory elements of the cold response in zebrafish. Nucleic Acids Research, 2015. 43(19): p. 9198–9213.

7. Star, B., et al., The genome sequence of Atlantic cod reveals a unique immune system. Nature, 2011. 477(7363): p. 207–210.

8. Chen, Z.Z., et al., Transcrintomic and genomic evolution under constant cold in Antarctic notothenioid fish. Proceedings of the National Academy of Sciences of the United States of America, 2008. 105(35): p. 12944–12949.

9. Fletcher, G.L., C.L. Hew, and P.L. Davies, Antifreeze proteins of teleost fishes. Annual Review of Physiology, 2001. 63: p. 359–390.

10. Verde, C., et al., The Hemoglobins of Fishes Living at Polar Latitudes - Current Knowledge on Structural Adaptations in a Changing Environment. Current Protein & Peptide Science, 2008. 9(6): p. 578–590.

11. Eastman, J.T. and M.J. Lannoo, Brain and sense organ anatomy and histology in hemoglobinless Antarctic icefishes (Perciformes : Notothenioidei : Channichthyidae). Journal of Morphology, 2004. 260(1): p. 117–140.

12. Parker, S.K. and H.W. Detrich, Evolution, organization, and expression of alpha-tubulin genes in the Antarctic fish Notothenia coriiceps - Adaptive expansion of a gene family by recent gene duplication, inversion, and divergence. Journal of Biological Chemistry, 1998. 273(51): p. 34358–34369.

13. Barnes, K.R., et al., Cold acclimation of NaCl secretion in a eurythermic teleost: Mitochondrial function and gill remodeling. Comparative Biochemistry and Physiology a-Molecular & Integrative Physiology, 2014. 168: p. 50–62.

14. Jayasundara, N., et al., Proteomic analysis of cardiac response to thermal acclimation in the eurythermal goby fish Gillichthys mirabilis. Journal of Experimental Biology, 2015. 218(9): p. 1359–1372.

15. Costas, B., et al., Different environmental temperatures affect amino acid metabolism in the eurytherm teleost Senegalese sole (Solea senegalensis Kaup, 1858) as indicated by changes in plasma metabolites. Amino Acids, 2012. 43(1): p. 327–335.

16. Alzaid, A., et al., Cold-induced changes in stress hormone and steroidogenic transcript levels in cunner (Tautogolabrus adspersus), a fish capable of metabolic depression. General and Comparative Endocrinology, 2015. 224: p. 126–135.

17. Little, A.G. and F. Seebacher, Thyroid hormone regulates muscle function during cold acclimation in zebrafish (Danio rerio). Journal of Experimental Biology, 2013. 216(18): p. 3514–3521.

18. Little, A.G., et al., Thyroid hormone actions are temperature-specific and regulate thermal acclimation in zebrafish (Danio rerio). Bmc Biology, 2013. 11: p. 15.

19. Little, A.G. and F. Seebacher, Thyroid hormone regulates cardiac performance during cold acclimation in zebrafish (Danio rerio). J Exp Biol, 2014. 217(Pt 5): p. 718–25.

20. Roy, R., D. Ghosh, and A.B. Das, Homeoviscous adaptation of different memberanes in the brain of an air-breathing indian teleost, *channa-punctatus*, during seasonal variation of enviromental-temperature. Journal of Thermal Biology, 1992. 17(4-5): p. 209–215.

21. Logue, J.A., et al., Lipid compositional correlates of temperature-adaptive interspecific differences in membrane physical structure. Journal of Experimental Biology, 2000. 203(14): p. 2105–2115.

22. Iglesias, T.L., et al., Life in the unthinking depths: energetic constraints on encephalization in marine fishes. Journal of Evolutionary Biology, 2015. 28(5): p. 1080–1090.

23. Johnston, I.A., A. Clarke, and P. Ward, Temperature and Metabolic-Rate in Sedentary Fish from the Antarctic, North-Sea and Indo-West Pacific-Ocean. Marine Biology, 1991. 109(2): p. 191–195.

24. Clarke, A. and N.M. Johnston, Scaling of metabolic rate with body mass and temperature in teleost fish. Journal of Animal Ecology, 1999. 68(5): p. 893–905.

25. Martin-Perez, M., et al., New Insights into Fish Swimming: A Proteomic and Isotopic Approach in Gilthead Sea Bream. Journal of Proteome Research, 2012. 11(7): p. 3533–3547.

26. Speers-Roesch, B. and J.S. Ballantyne, Activities of antioxidant enzymes and cytochrome c oxidase in liver. of Arctic and temperate teleosts. Comparative Biochemistry and Physiology a-Molecular & Integrative Physiology, 2005. 140(4): p. 487–494.

27. Jayasundara, N., L.D. Gardner, and B.A. Block, Effects of temperature acclimation on Pacific bluefin tuna (Thunnus orientalis) cardiac transcriptome. American Journal of Physiology-Regulatory Integrative and Comparative Physiology, 2013. 305(9): p. R1010–R1020.

28. Mininni, A.N., et al., Liver transcriptome analysis in gilthead sea bream upon exposure to low temperature. Bmc Genomics, 2014. 15.

29. Chen, Y.X., et al., The Grass Carp Genome Database (GCGD): an online platform for genome features and annotations. Database-the Journal of Biological Databases and Curation, 2017: p. 8.

30. Johnston, I.A. and J. Dunn, Temperature acclimation and metabolism in ectotherms with particular reference to teleost fish. Symposia of the Society for Experimental Biology, 1987. 41: p. 67–93.

31. Crawshaw, L.I., Low-temperature dormancy in fish. American Journal of Physiology, 1984. 246(4): p. R479–R486.

32. Nelson, E.R. and H.R. Habibi, Estrogen receptor function and regulation in fish and other vertebrates. General and Comparative Endocrinology, 2013. 192: p. 15–24.

33. Hernandez, I., et al., Effect of seasonal acclimatization on estrogen-induced vitellogenesis and on the hepatic estrogen receptors in the male carp. Biochemistry International, 1992. 28(3): p. 559–567.

34. Verderame, M. and R. Scudiero, A comparative review on estrogen receptors in the reproductive male tract of non mammalian vertebrates. Steroids, 2018. 134: p. 1–8.

35. Wade, G.N., J.E. Schneider, and H.Y. Li, Control of fertility by metabolic cues. American Journal of Physiology-Endocrinology and Metabolism, 1996. 270(1): p. E1–E19.

36. Ernst, R., C.S. Ejsing, and B. Antonny, Homeoviscous Adaptation and the Regulation of Membrane Lipids. Journal of Molecular Biology, 2016. 428(24): p. 4776–4791.

37. G. J., Thompson,. Mechanisms of homeoviscous adaptation in membranes. Cellular Acclimatisation to Environmental Change, 1983.

38. Gracey, A.Y., et al., Coping with cold: An integrative, multitissue analysis of the transcriptome of a poikilothermic vertebrate. Proceedings of the National Academy of Sciences of the United States of America, 2004. 101(48): p. 16970–16975.

39. Xu, D., et al., Transcriptional response to low temperature in the yellow drum (Nibea albiflora) and identification of genes related to cold stress. Comparative Biochemistry and Physiology D-Genomics & Proteomics, 2018. 28: p. 80–89.

40. Iida, K., et al., Genome-wide analysis of alternative pre-mRNA splicing in Arabidopsis thaliana based on full-length cDNA sequences. Nucleic Acids Research, 2004. 32(17): p. 5096–5103.

41. Calixto, C.P.G., et al., Rapid and Dynamic Alternative Splicing Impacts the Arabidopsis Cold Response Transcriptome. Plant Cell, 2018. 30(7): p. 1424–1444.

42. Polley, S.D., et al., Differential expression of cold- and diet-specific genes encoding two carp liver Delta 9-acyl-CoA desaturase isoforms. American Journal of Physiology-Regulatory Integrative and Comparative Physiology, 2003. 284(1): p. R41–R50.

43. Long, Y., et al., Transcriptomic Characterization of Temperature Stress Responses in Larval Zebrafish. Plos One, 2012. 7(5).

44. Healy, T.M. and P.M. Schulte, Patterns of alternative splicing in response to cold acclimation in fish. J Exp Biol, 2019.

45. Teige, M., et al., The MKK2 pathway mediates cold and salt stress signaling in Arabidopsis. Molecular Cell, 2004. 15(1): p. 141–152.

46. Chinnusamy, V., K. Schumaker, and J.K. Zhu, Molecular genetic perspectives on cross-talk and specificity in abiotic stress signalling in plants. Journal of Experimental Botany, 2004. 55(395): p. 225–236.

47. Wen, X., et al., iTRAQ-based quantitative proteomic analysis of Takifugu fasciatus liver in response to low-temperature stress. Journal of Proteomics, 2019. 201: p. 27–36.

48. Niu, H., et al., The role of dusp1 downregulation in apoptosis of zebrafish ZF4 cells under cold stress. Journal of Fishery Sciences of China, 2017. 24(5): p. 995–1002.

49. Cai, J.-H., et al., Temperature-induced embryonic diapause in blue-breasted quail (Coturnix chinensis) correlates with decreased mitochondrial-respiratory network and increased stress-response network. Poultry science, 2019. 98(7): p. 2977–2988.

50. Kanzleiter, T., et al., Evidence for Nr4a1 as a cold-induced effector of brown fat thermogenesis. Physiological Genomics, 2005. 24(1): p. 37–44.

51. Malki, K., et al., Transcriptome analysis of genes and gene networks involved in aggressive behavior in mouse and zebrafish. American Journal of Medical Genetics Part B-Neuropsychiatric Genetics, 2016. 171(6): p. 827–838.

52. Lopes, J.S., R. Abril-de-Abreu, and R.F. Oliveira, Brain Transcriptomic Response to Social Eavesdropping in Zebrafish (Danio rerio). Plos One, 2015. 10(12).

53. Dietz, T.J. and G.N. Somero, Species-specific and tissue-specific synthesis patterns for heat-shock proteins Hsp70 and Hsp90 in several marine teleost fishes. Physiological Zoology, 1993. 66(6): p. 863–880.

54. Iwama, G.K., et al., Heat shock proteins and physiological stress in fish. American Zoologist, 1999. 39(6): p. 901–909.

55. Gantner, M.L., et al., GADD4 gamma regulates the thermogenic capacity of brown adipose tissue. Proceedings of the National Academy of Sciences of the United States of America, 2014. 111(32): p. 11870–11875.

56. Patel, R.K. and M. Jain, NGS QC Toolkit: A Toolkit for Quality Control of Next Generation Sequencing Data. Plos One, 2012. 7(2).

57. Trapnell, C., et al., Differential gene and transcript expression analysis of RNA-seq experiments with TopHat and Cufflinks. Nature Protocols, 2012. 7(3): p. 562–578.

58. Pertea, M., et al., StringTie enables improved reconstruction of a transcriptome from RNA-seq reads. Nature Biotechnology, 2015. 33(3): p. 290-+.

59. Kang, Y.J., et al., CPC2: a fast and accurate coding potential calculator based on sequence intrinsic features. Nucleic Acids Research, 2017. 45(W1): p. W12–W16.

60. Wang, L., et al., CPAT: Coding-Potential Assessment Tool using an alignment-free logistic regression model. Nucleic Acids Research, 2013. 41(6).

61. Patro, R., et al., Salmon provides fast and bias-aware quantification of transcript expression. Nature Methods, 2017. 14(4): p. 417-+.

62. Robinson, M.D., D.J. McCarthy, and G.K. Smyth, edgeR: a Bioconductor package for differential expression analysis of digital gene expression data. Bioinformatics, 2010. 26(1): p. 139–140.

63. Lun, A.T., Y. Chen, and G.K. Smyth, It’s DE-licious: A Recipe for Differential Expression Analyses of RNA-seq Experiments Using Quasi-Likelihood Methods in edgeR. Methods Mol Biol, 2016. 1418: p. 391–416.

